# YAP1 is essential for self-organized differentiation of pluripotent stem cells

**DOI:** 10.1101/2022.09.29.510043

**Authors:** Kira Zeevaert, Roman Goetzke, Mohamed H. Elsafi Mabrouk, Marco Schmidt, Catharina Maaßen, Ann-Christine Henneke, Chao He, Arnold Gillner, Martin Zenke, Wolfgang Wagner

## Abstract

The Yes-associated protein 1 (YAP1) is a downstream effector of the Hippo pathway and essential mechanotransducer. It has been suggested to play a crucial role for early embryo development, but the relevance for early germ layer commitment of human induced pluripotent stem cells (iPSCs) remains largely unclear. To gain better insight into the function of YAP1 in these early cell-fate decisions, we generated iPSC lines with YAP1 knockout (*YAP*^-/-^) with CRISPR/Cas9 technology and analyzed transcriptomic and epigenetic modifications. In *YAP*^-/-^ iPSCs the expression of several YAP1 targets changed and *NODAL*, which is an important regulator of cell differentiation, was upregulated. Furthermore, YAP1 deficiency evoked global DNA methylation changes. Directed differentiation of adherent iPSC colonies toward endoderm, mesoderm, and ectoderm could be induced, albeit endodermal and ectodermal differentiation showed transcriptomic and epigenetic changes in *YAP*^-/-^ lines. Notably, in self-organized embryoid bodies (EBs) germ layer specification was clearly impaired. This phenotype was rescued via lentiviral overexpression of *YAP1* and in tendency also by NODAL inhibitors. Our results demonstrate that YAP1 plays an important role during early germ layer specification of iPSCs, particularly for the non-directed self-organization of EBs, and this is at least partly attributed to activation of the NODAL pathway.

## Introduction

Embryoid bodies (EBs) resemble self-organized three-dimensional (3D) aggregates of induced pluripotent stem cells (iPSCs) (Poh et al., 2014; Zeevaert et al., 2020). They recapitulate some aspects of blastocyst formation and gastrulation after fertilization (Doetschman et al., 1985), and give rise to all three germ layers – endoderm, mesoderm, and ectoderm (Itskovitz-Eldor et al., 2000; Montero and Heisenberg, 2004). Representative markers for germ layer specification include GATA-binding factor 6 (*GATA6)* for endoderm, t-box transcription factor T (*TBXT*, brachyury) for mesoderm, and paired box protein pax-6 (*PAX6)* for ectoderm (Gifford et al., 2013; Sheridan et al., 2012). Cell-fate decisions are reflected by epigenetic changes, including DNA methylation (DNAm) changes that play a crucial role for regulation of gene expression and chromatin organization during mammalian development (Passaro et al., 2021; Smith and Meissner, 2013). However, it is largely unclear which signaling cascades are involved in the self-organized differentiation of EBs.

*In vivo*, tissue formation is highly dependent from the spatiotemporal regulation of different signaling pathways, such as the WNT pathway or the transforming growth factor beta (TGF-β) pathway, with one of its main ligands being NODAL (Gadue et al., 2006; Shen, 2007; Wu and Hill, 2009). Furthermore, NODAL seems to play a central role for the cellular organization within iPSC colonies (Tewary et al., 2017; Warmflash et al., 2014). Cells at the outer rim of colonies upregulate pluripotency-associated genes as well as the TGF-β pathway, particularly NODAL and its inhibitor left-right determination factor 1 (LEFTY1) (Elsafi Mabrouk et al., 2022). Another important signaling pathway of early embryo development is the Hippo pathway (Wu and Guan, 2021). Once activated, the Hippo pathway limits tissue growth and cell proliferation and is suggested to affect growth of embryonic structures (Low et al., 2014; Moya and Halder, 2019; Nishioka et al., 2009). Through a cascade of phosphorylation events the Hippo pathway ultimately results in phosphorylation and degradation of the Yes-associated protein 1 (YAP1). In contrast, activated YAP1 binds to the TEA domain family member 1-4 DNA-binding (TEAD) proteins and regulates transcription of a variety of genes (Moya and Halder, 2019).

Cellular differentiation is not only guided by soluble signals, but also by mechanical cues. In fact, YAP1 is also an essential mechanotransducer and translocates to the nucleus in response to substrate elasticity and cell shape, independent from the Hippo cascade (Aragona et al., 2013; Dupont et al., 2011). There is accumulating evidence that YAP1 transfers mechanical signals during germ layer formation in mammalian embryos (Pagliari et al., 2021; Wu and Guan, 2021). YAP1 deficiency causes mouse embryonic developmental arrest around E8.5. with defective yolk sac vasculogenesis, chorioallantoic attachment, and embryonic axis elongation (Morin-Kensicki et al., 2006). YAP1 is also important for germ cell differentiation from mouse epiblast stem cells (Kagiwada et al., 2021). Furthermore, YAP1 is required to maintain human pluripotent stem cells in naïve state (Qin et al., 2016) and increased YAP1 activity supports tissue regeneration (Elster and von Eyss, 2020). In human embryonic stem cells (hESCs) YAP1 represses mesendodermal differentiation by regulating the activity of the NODAL and WNT3 pathway (Estarás et al., 2017; Stronati et al., 2022). It can therefore be anticipated that YAP1 is also relevant for the self-organization and cell-fate decisions in EBs, but this aspect has so far not been systematically addressed.

In this study, we generated human iPSCs with YAP1 knockout (*YAP*^*-/-*^) to further explore transcriptomic and epigenetic changes at pluripotent state and upon directed differentiation towards mesoderm, endoderm, and ectoderm. Overall, *YAP*^*-/-*^ iPSCs revealed changes in gene expression and DNAm after endodermal and ectodermal differentiation, but they remained capable of trilineage differentiation. In contrast, in self-organized EBs, germ layer specification was clearly impaired – *GATA6, TBXT*, and *PAX6* were hardly up-regulated. This phenotype was rescued via lentiviral overexpression of *YAP1* and in tendency also by NODAL inhibitors. Our results indicate that YAP1 is essential for early germ layer specification of iPSCs, particularly for the non-directed self-organization of EBs, and this is attributed to activation of the NODAL pathway.

## Experimental Procedures

### Cell culture and directed differentiation

Three human iPSC lines were generated from bone marrow derived mesenchymal stromal cells – hPSCreg: UKAi009-A (wildtype (WT) 1), UKAi010-A (WT 2), and UKAi011-A (WT3) – by reprogramming with episomal plasmids (Goetzke et al., 2018). All samples were taken after informed and written consent using guidelines approved by the Ethic Committee for the Use of Human Subjects at the University of Aachen (permit number: EK128/09). All methods were performed in accordance with the relevant guidelines and regulations. The iPSC lines were cultured on tissue culture plastic coated with vitronectin (0.5 μg/cm^2^; Stemcell Technologies, Vancouver, Canada) in StemMACS iPS-Brew XF (Miltenyi Biotec GmbH, Bergisch Gladbach, Germany). Pluripotency was validated by three lineage differentiation potential and Epi-Pluri-Score analysis, as described in our previous work (Lenz et al., 2015).

For directed differentiation towards endodermal, mesodermal, and ectodermal lineage we used the STEMdiff Trilineage Differentiation Kit (Stemcell Technologies, Vancouver, Canada), according to the manufacturer’s instructions. Endoderm and mesoderm differentiation was performed for 5 days, while ectoderm differentiation lasted 7 days. As an undifferentiated control, iPSCs were kept in StemMACS iPS-Brew XF for 5 days.

### CRISPR/Cas9 mediated YAP1 knockout

Two YAP1 knockout (*YAP*^*-/-*^) cell lines (iPSC *YAP*^*-/-*^ *1* and iPSC *YAP*^*-/-*^ *2*) were generated form WT 1 with a CRISPR/Cas9 nuclease approach. One guide RNA was designed to target exon three of the human *YAP1* gene, which is shared in all eleven transcripts. Cutting the DNA strand in exon three led to non-homologous end joining which goes along with insertion-deletion mutations. Reading frameshifts generated a complete knockout of the YAP1 protein. For ribonucleoprotein (RNP) delivery into iPSCs Alt-R CRISPR-Cas9 crRNA, Alt-R CRISPR-Cas9 tracrRNA, and Alt-R HiFi Sp. Cas9 Nuclease (all IDT, Coralville, USA) were used. After RNP assembly, iPSC were transfected using the NEON transfection system (1300 V, 30 ms pulse width, 1 pulse, Thermo Fischer Scientific, Waltham, USA). Electroporated cells were seeded in StemMACS iPS-Brew XF with Rho-associated protein kinase (ROCK) inhibitor Y-27632 (10 μM, Abcam, Cambridge, UK) and 1x CloneR (Stemcell Technologies, Vancouver, Canada) on laminin 521 (5 μg/mL, BioLamina, Sundbyberg, Sweden) coated plates. Ten days after transfection, cells were singularized and seeded on mouse embryonic fibroblasts (MEFs). Eight days later, individual colonies were picked and expanded on feeder-free vitronectin-coated plates. Expanded clones were screened for homozygous mutations in exon three via Sanger sequencing (Eurofins Genomics, Ebersberg, Germany; sequencing primer: GAGCAGTGAGATGCTGTGAC).

### *YAP1* overexpression

Lentiviruses with pGAMA-YAP (plasmid #74942, Addgene, Watertown, USA) (Qin et al., 2016) were produced in HEK-293T cells. 8 μg/mL Polybrene were added to virus suspension and incubated for 30 minutes at 37°C. For infection, iPSCs were centrifuged with virus-Polybrene-StemMACS iPS-Brew XF solution for 2 hours at 1,500 rpm at room temperature (RT) and incubated at 37°C for 1 hour. After 24 hours of culture in StemMACS iPS-Brew XF the infection of iPSCs was repeated with an incubation of 3 hours at 37°C. Ten days after infection, cells were singularized and seeded on MEFs. Individual colonies were then picked based on fluorescent mCherry labeling of the pGAMA-YAP plasmid and expanded for one passage on MEFs and after that on vitronectin-coated plates.

### iPSCs on sub-μm surface topography

Structured polyimide (PI) foils were generated in continuation of our previous work (Abagnale et al., 2015), by multi-beam interference technology using a UV laser with 38 ns pulse duration, 300 μm spot size, and 33 % overlap. Samples were imaged with a scanning electronic microscope (SEM) LEO 1455 EP (Carl Zeiss, Jena, Germany) as described before (Abagnale et al., 2015). PI was coated with vitronectin and single iPSCs were seeded with 10,000 cells/cm^2^ in StemMACS iPS-Brew XF with rho-associated protein kinase (ROCK) inhibitor Y-27632 (10 μM). Elongation of single iPS cells (day 1 after seeding) and iPSC colonies (day 2 after seeding) on linear periodic groove-ridge structures with a periodicity of 665 nm and a depth of 200 nm was imaged with an EVOS FL (Thermo Fisher, Waltham, USA) at 4x magnification. Aspect ratios (length/diameter) were quantified using ImageJ.

### Spatially confined iPSC colonies and embryoid body formation

Micro-contact printing (μCP) of vitronectin was carried out to generate circular adhesion islands with a diameter of 600 μm that facilitate attachment of iPSCs (Elsafi Mabrouk et al., 2022). The iPSC lines were seeded with 10,000 cells/cm^2^ in StemMACS iPS-Brew XF with ROCK inhibitor Y-27632 (10 μM) added for the first day. The iPSC growth was restricted to the μCP area with upregulation of pluripotency associated markers at the rim (Elsafi Mabrouk et al., 2022). After six to eight days, the iPSC colonies self-detached spontaneously from the vitronectin-μCP substrates (Elsafi Mabrouk et al., 2022). When more than 50 % of the colonies detached, aggregates were harvested and considered as day 0 for further differentiation steps. Multilineage differentiation of EBs was performed with differentiation induction medium (EB-medium) containing Knockout DMEM (Gibco, Carlsbad, USA), 20 % v/v FCS (Lonza, Basel, Switzerland), 2 mM L-Glutamine, 1x Non-Essential amino acids, 100 nM b-Mercaptoethanol, and 100 U/mL Penicillin/Streptomycin solution. EBs were differentiated in ultra-low attachment plates (Corning, New York, USA) for 5 days. Medium was changed every second day. For inhibition of NODAL signaling, the EB-medium was supplemented with the TGF-β-Type I-receptor inhibitors SB-431542 (10 μM; MedChemExpress, Monmouth Junction, USA) (Inman et al., 2002) or A83-01 (100 nM; Stemcell Technologies, Vancouver, Canada) (Tojo et al., 2005) for 5 days with medium changes every second day.

### Western Blot

Cells were harvested and total cell extracts were prepared in cold lysis buffer (RIPA buffer, 10 mM NaF, 1mM Na3VO4, and 7x complete mini protease inhibitor). Protein concentration was determined by Bradford protein assay, measured in a photometer (Infinite 200 PRO, Tecan Trading AG, Switzerland). 20 μg protein per sample were incubated with 4x SDS Protein Sample Buffer (50 mM Tris/HCl pH 6.8, 2 % SDS, 0.01 % Bromophenol blue, 2.5 % β-mercapthoethanole and 10 % glycerol) for 5 minutes at 99°C. Samples were separated in 12 % Mini-PROTEAN TGX Precast Protein Gels (Bio-Rad, München, Germany) and transferred onto polyvinylidene fluoride (PVDF; Merck Millipore, Burlington, USA). Membrane was blocked for 1 hour at RT in 4 % w/v bovine serum albumin (BSA) and incubated with primary antibodies against YAP1 and actin (Supplemental Table 1) in blocking solution overnight at 4°C. Secondary antibodies (Supplemental Table 1) were incubated for 1 hour at RT and protein bands were visualized using a ChemiDoc XRS+ (Bio-Rad Laboratories, Hercules, USA).

### Immunostaining

Cells were fixed with 4 % paraformaldehyde (PFA) for 20 minutes treated with PBS containing 1 % w/v BSA and 0.1 % v/v Triton X-100 (Bio-Rad, München, Germany) for 30 minutes, and then incubated overnight at 4°C with primary antibodies against YAP1, WW domain-containing transcription regulator protein1 (WWTR1, TAZ), octamer-binding transcription factor 4 (OCT4, POU5F1), GATA6, brachyury, PAX6, and NODAL (Supplemental Table 1). Secondary antibody (Supplemental Table 1) staining was done at RT for 1 hour. Samples were counterstained with DAPI (10 ng/mL) for 15 minutes at RT in the dark. 2D samples were imaged using either a LSM 700 confocal microscopy (Carl Zeiss, Wetzlar, Germany) using 20x objective or EVOS FL (Thermo Fisher, Waltham, USA) at 4x, 10x, and 20x magnification or with an Axioplan 2 fluorescence microscope (Thermo Fisher, Waltham, USA) at 20x magnification. Radial profile of immunofluorescence images of 2D μCP colonies were quantified using radial profile extended ImageJ plugin.

For staining of 3D aggregates, EBs were fixed with 4 % PFA for 1 hour. Thereafter, fixed EBs were embedded in fibrin gels which consisted of fibrinogen (5 mg/mL), CaCl2 (3.75 mM), and thrombin (20 U/mL); all fibrin gel components were dissolved in tris-buffered saline (TBS)-buffer composed of Tris-Cl (50 mM), NaCl (150 mM), and deionized water (pH 7.5; all Sigma Aldrich, Hamburg, Germany). Blocking and permeabilization was done in PBS with 1 % w/v BSA and 0.1 % v/v Triton-X 100 for 1 hour at RT. Samples were stained with primary antibodies against YAP1, OCT4, GATA6, brachyury, and PAX6 (Supplemental Table 1) at 4°C overnight. Secondary antibody (Supplemental Table 1) staining was performed at 4°C overnight. Finally, aggregates were counterstained with DAPI (1 ng/mL) for 2 more days at 4°C. EBs were imaged using FV1000MPE two-photon microscope (Olympus Corp., Tokyo, Japan) using 35x water-immersion objective.

### Transcriptomic analysis

RNA was isolated from undifferentiated iPSCs, differentiated iPSCs, and EBs at day 5 using the NucleoSpin RNA Plus Kit (Macherey-Nagel, Düren, Germany) and quantified with a NanoDrop ND-2000 spectrophotometer (Thermo Scientific, Wilmington, USA). Library preparation (QuantSeq 3’-mRNA) and RNA-sequencing were performed by Life & Brain company (Bonn, Germany) on a NovaSeq 6000 sequencer (100 bp/read). Quality of FASTAQ files was quantified using FastQC, and adaptor sequences and low quality reads were trimmed using Trimmomatic. Alignment of the reads was done using STAR (hg38 genome build) and the resulting count matrices were normalized with the variance-stabilized transformation (VST) method and principal component analysis of the normalized and transformed data were done using the DESeq2 package in R (Love et al., 2014). Differential gene expression analysis between YAP1 knockout and wildtype samples was performed using Wald-test and p-values were adjusted with the Benjamini-Hochberg procedure. Genes with adjusted p-value < 0.05 and fold change > 2 were considered as differentially expressed. Pathway activation was inferred with the PROGENy package (Schubert et al., 2018) using top 100 pathway responsive genes. To get p-values for the differentially activated pathways, resulting Z-scores were transformed to p-values using the pnorm function from stats R package and adjusted using the Benjamini-Hochberg procedure. Marker genes of germ layer differentiation and stem cells were identified from previously published marker datasets (Maguire et al., 2013; Stronati et al., 2022). Gene set enrichment analysis (GSEA) was carried out for a fold-change ranked gene list using the biological process section of the gene ontology database and the fgsea algorithm implemented by ClusterProfiler R package (Wu et al., 2021).

### DNA methylation profiling

Genomic DNA was isolated from iPSCs and EBs using the NucleoSpin Tissue Kit (Macherey-Nagel, Düren, Germany) with the manufacturer’s instructions and quantified with a NanoDrop ND-2000 spectrophotometer (Thermo Scientific, Waltham, USA). 1.2 μg DNA was bisulfite converted and analyzed with Illumina Beadchip Epic Array at Life & Brain (Bonn, Germany). Raw IDAT files were used for quality control and ssNoob normalization (Triche et al., 2013) with the minfi R package (Aryee et al., 2014). Detection p-values were calculated with the SeSAMe R package and the pOOBAH approach. CpG sites on XY chromosomes, non-cg probes, probes with a detection p-value > 0.05 in two or more samples, SeSAMes masked probes (Zhou et al., 2017; Zhou et al., 2018), and probes flagged in Illumina epic manifest version b5 were removed. This reduced the number to 645,055 CpG sites, which were used for further analysis. The Limma R package was used for generating the multidimensional scaling (MDS) plots and calculating BH adjusted p-values for differentially methylated sites accounting for sample donors. CpG sites with a difference of mean beta values ≥ 0.1 and an adjusted p-value ≤ 0.05 were considered significantly different. To select the CpGs, which change during EB differentiation, the days (0, 4, and 7) were considered as covariate in the Limma design matrix. The Epi-Pluri-Score was calculated as described before (Lenz et al., 2015). For combining DNAm and gene expression data we used the Illumina BeadChip annotation and merged the data by matching to Ensembl IDs, only considering CpGs in promoter regions (TSS1500 and TSS200). The R packages ggplot2 and ComplexHeatmap were used for generating the figures.

### Semi-quantitative reverse-transcriptase PCR

Total RNA was isolated from iPSCs and EBs using the NucleoSpin RNA Plus Kit (Macherey-Nagel, Düren, Germay), quantified with a NanoDrop ND-2000 spectrophotometer (Thermo Scientific, Waltham, USA) and converted into cDNA using the High-Capacity cDNA Reverse Transcription Kit (Applied Biosystems, Waltham, USA). Semi-quantitative reverse-transcriptase PCR (RT-qPCR) was either carried out using TaqMan gene expression master mix and gene-specific TaqMan assays or Power SYBR Green PCR Master Mix (Applied Biosystems, Waltham, USA) and gene-specific primers in a StepOnePlus machine (Applied Biosystems, Waltham, USA). Primers for *POU5F1, GATA6, TBXT, PAX6*, and housekeeping gene glyceraldehyde 3-phosphate dehydrogenase (*GAPDH)* are provided in Supplemental Table 2. TaqMan assays for *YAP1* and *GAPDH* are provided in Supplemental Table 3.

## Results

### YAP1 deficiency induces transcriptomic and epigenetic modulation in human iPSCs

We generated two YAP1 knockout iPSC lines (*YAP*^*-/-*^ 1 and *YAP*^*-/-*^ 2) with CRISPR/Cas9 technology by homozygous induction of reading frame shift mutations in exon three of the human *YAP1* gene (Suppl. Figure 1A). YAP1 deficiency was confirmed via Western Blot analysis (Figure 1A) and immunocytochemistry (Figure 1B, Suppl. Figure 1B). The pluripotent state was not impaired by YAP1 knockout, since there was no obvious impact on iPSC colony morphology or OCT4 expression (Figure 1B, Suppl. Figure 1B). Furthermore, the Epi-Pluri-Score (Lenz et al., 2015) remained positive in *YAP*^*-/-*^ lines (Suppl. Figure 1C). Despite these phenotypic similarities, there were significant differences between the transcriptomes of wildtype (n = 3) and knockout (n = 2) iPSC lines. RNA sequencing revealed 100 genes that were significantly downregulated and 91 genes that were upregulated in *YAP*^*-/-*^ iPSCs (log_2_ fold change > 1; adj. p-value < 0.05; Figure 1C). When we compared the differentially expressed genes with 2,511 YAP1 targets, including long non-coding RNAs (lncRNA) and protein coding regions without intergenic regions, from a public available YAP1 chromatin immunoprecipitation DNA-sequencing (ChIP-Seq) dataset (Pagliari et al., 2021), we observed a significant overlap (Fisher exact test: adj. p-value = 0.035; Odds ratio = 1.48), indicating that the transcriptomic changes are directly modulated by YAP1 deficiency (Figure 1D). Interestingly, mesoderm related YAP1 target genes, such as hes related family BHLH transcription factor with YRPW motif 2 (*HEY2*) and RNA binding motif protein 47 (*RBM47*) were upregulated in YAP1 deficient iPSCs compared to wildtype iPSCs (Figure 1D), supporting previous observations that mesodermal differentiation might be improved in YAP1 knockout cells (Estarás et al., 2017; Stronati et al., 2022). Furthermore, downregulation of gene expression was significantly enriched in the WNT pathway (adj. p-value = 0.0026), whereas *NODAL*, which is part of the TGF-β superfamily and essential for the regulation of pluripotency maintenance and embryogenesis, was significantly upregulated in *YAP*^*-/-*^ iPSCs (adj. p-value = 6.04*10^−7^, log_2_ fold change = 6.06).

**Figure 1:**
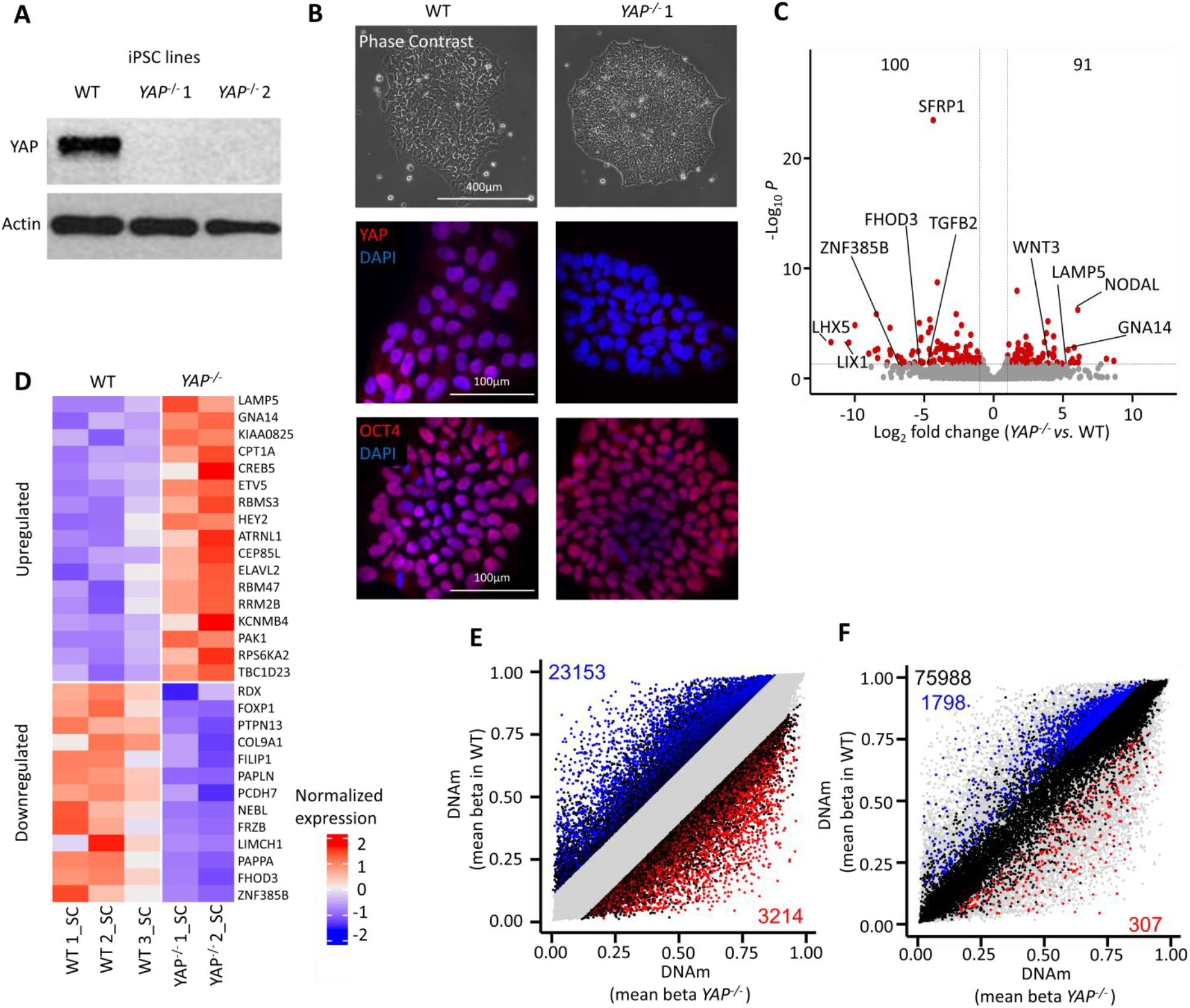
YAP1 deficiency induces transcriptomic and epigenetic modulation in human iPSCs. **(A)** Western Blot analysis of wildtype (WT) and *YAP*^*-/-*^ iPSC lines with antibodies against YAP1 (65 kD) and the housekeeping protein actin (42 kD). **(B)** Morphologic and immunofluorescence analysis of WT and *YAP*^*-/-*^ 1 iPSC colonies with antibodies against YAP1 and OCT4 (nuclei counterstained with DAPI). **(C)** RNA sequencing analysis of *YAP*^*-/-*^ (n = 2) and WT (n = 3) iPSCs. 100 genes are significantly downregulated and 91 genes are significantly upregulated in *YAP*^*-/-*^ (log_2_ fold change > 1; adj. p-value < 0.05). Specific YAP1 target genes and genes related to TGF-β or WNT pathway, as well as germ layer related genes are highlighted. **(D)** Normalized expression of 17 up- and 13 downregulated genes which are significantly differentially expressed between WT and *YAP*^*-/-*^ iPSCs and which are YAP1 targets according to human YAP1 ChIP-Seq data (Pagliari et al., 2021). **(E)** YAP1 deficiency evokes global DNA methylation changes between *YAP*^*-/-*^ (n = 2) and WT (n = 3) iPSCs, measured via Illumina Beadchip Epic Array (beta values ranging from 0 to 1). 23,153 CpGs are significantly hypomethylated and 3,214 CpGs are significantly hypermethylated in *YAP*^*-/-*^ iPSCs (adj. p-value ≤ 0.05; difference in mean ≥ 0.1). **(F)** 78,093 CpGs are related to 2,511 YAP1 target genes (Pagliari et al., 2021). Only 1,798 of these CpGs are significantly hypomethylated, and 307 CpGs significantly hypermethylated between *YAP*^*-/-*^ and WT iPSCs indicating that DNAm changes are not clearly enriched in YAP1 target genes.

To further explore if the YAP1 deficiency also entailed epigenetic changes in iPSCs, we compared DNAm profiles of *YAP*^*-/-*^ (n = 2) *versus* wildtype (n = 3) iPSCs (Figure 1E). 23,153 CG dinucleotides (CpG sites) revealed significant hypomethylation, whereas 3,214 CpGs gained DNAm in the *YAP*^*-/-*^ lines (adj. p-value ≤ 0.05; difference in mean ≥ 0.1). To check if these epigenetic modifications were also directly governed by the absence of YAP1 binding to specific sites in the genome we determined the overlap with CpGs in previously used YAP1 target genes (Pagliari et al., 2021). We identified 78,093 CpGs in our data set that are annotated to the 2,511 YAP1 target genes from which 1,798 CpGs are significantly hypomethylated and 307 significantly hypermethylated in the comparison of *YAP*^*-/-*^ *versus* wildtype iPSCs (Figure 1F). Overall, there is no clear enrichment of these significant differently methylated sites in genes, which were identified as YAP1 targets (Fisher exact test: adj. p-value = 1). Yet, we observed that in tendency genes with hypermethylation in promoter regions were downregulated on gene expression level, and *vice versa* (Suppl. Figure 1D). Taken together, already at non-differentiated stage, the *YAP*^*-/-*^ lines revealed transcriptomic and epigenetic differences.

### *YAP*^*-/-*^ iPSC colonies self-organize in 2D

Since YAP1 is an important mechanotransducer, and thereby relevant for biomaterial interaction, we furthermore analyzed the impact of YAP1 deficiency on sensing and processing mechanobiological stimulation. We have previously demonstrated that YAP1 activity was modulated in iPSCs on sub-micrometer structures upon BMP4-stimulation (Abagnale et al., 2017). In continuation of this work, we have now seeded wildtype and *YAP*^*-/-*^ cells on structured and unstructured polyimide (PI) foils (Figure 2A). The groove-ridge structures, with a periodicity of 665 nm and a depth of 200 nm, induced elongation of single individual cells (Figure 2B) as well as small iPSC colonies (Figure 2C). Aspect ratio of single iPSCs and iPSC colonies was about 5-fold (p < 0.0001) increased on structured compared to unstructured PI. Surprisingly, *YAP*^*-/-*^ iPSCs showed similar elongation on structured PI as wildtype cells (Figure 2D), indicating that YAP1 does not play a central role for this response to substrate structures.

**Figure 2:**
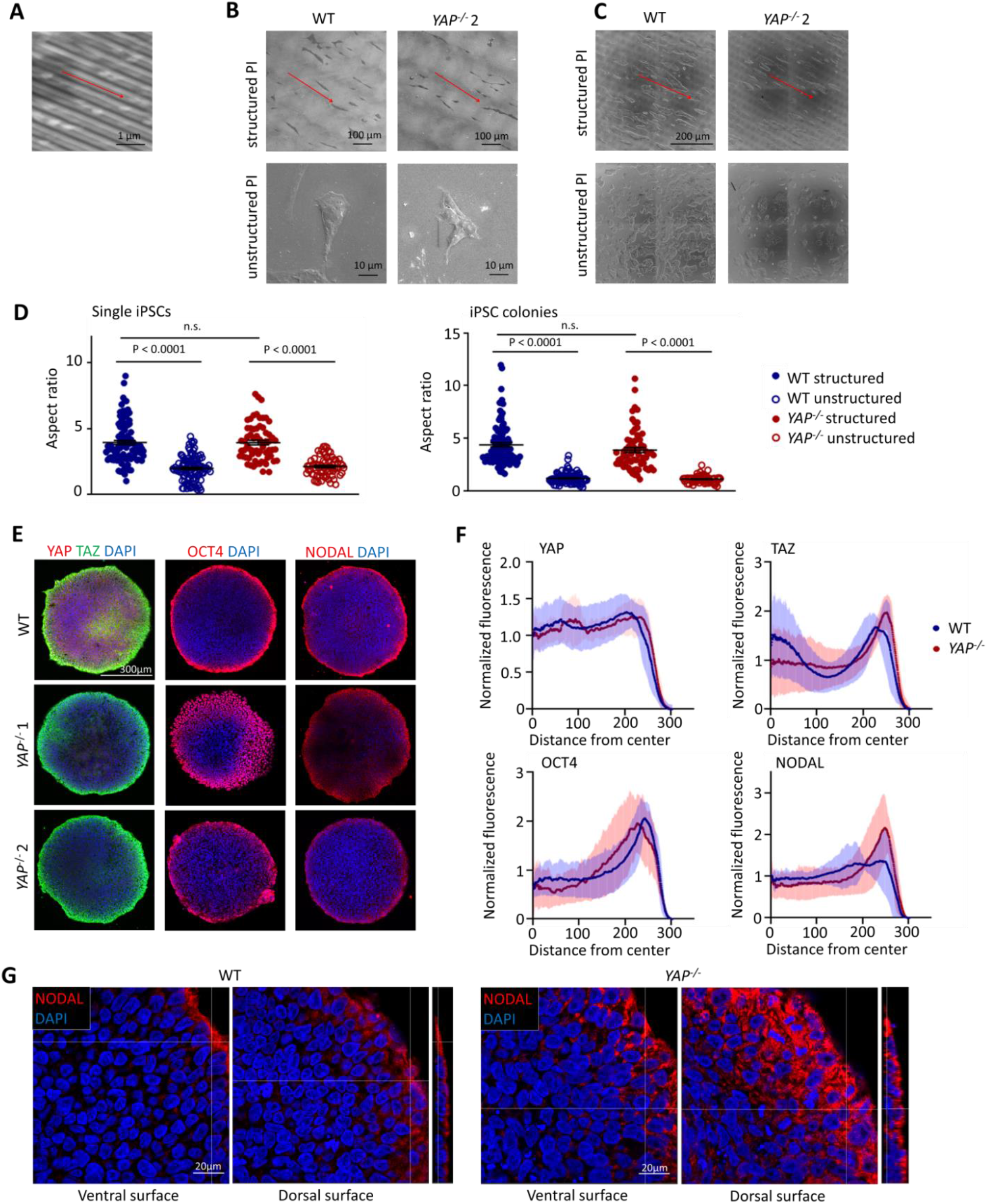
*YAP*^*-/-*^ iPSC colonies grow on sub-μm surface structures and self-organize in 2D. **(A)** Scanning electron microscopy image of structured polyimide (PI). PI was structured by multi-beam interference technology using a UV laser with 38 ns pulse duration, 300 μm spot size, and 33% overlap. Substrates have a linear periodic groove-ridge structure with a periodicity of 665 nm and a depth of 200 nm. **(B)** Scanning electron microscopy images of single WT and *YAP*^*-/-*^ iPSCs (day one after seeding) on structured and unstructured PI. **(C)** Light microscopy images of WT and *YAP*^*-/-*^ iPSC colonies (day two after seeding) on structured and unstructured PI. **(D)** Aspect ratios of WT (n = 3) and *YAP*^*-/-*^ (n = 2) on structured and unstructured PI at day one and day two after seeding were quantified with ImageJ. Unpaired t-test. P-values > 0.05 are considered as not significant (n.s.). **(E)** Confocal microscopy images of self-organized iPSC colonies of WT and *YAP*^*-/-*^ iPSCs at day 6. Geometrically confined colonies were stained with antibodies against YAP1, TAZ, OCT4, and NODAL (nuclei counterstained with DAPI). **(F)** Quantification of immunofluorescence signals of WT and *YAP*^*-/-*^ iPSC colonies on micro-contact printed substrates (mean ± SD; n = 3 per marker). **(G)** Confocal microscopy images of NODAL expression at the ventral and dorsal surface and the section of the edge of confined colonies from WT and *YAP*^*-/-*^ iPSCs at day 6 (nuclei counterstained with DAPI).

Processing mechanical signals not only from substrate topography but from surrounding cells can induce self-organization and cellular differentiation (Mao et al., 2016). In fact, guiding heterogeneity and spatial disorder of pluripotent iPSC colonies by geometric confinement supports their self-organized patterning (Warmflash et al., 2014). We have recently demonstrated that particularly *NODAL* and its inhibitor *LEFTY* were amongst the most highly upregulated genes at the rim of spatially confined colonies, which might be modulated by YAP1 activity (Elsafi Mabrouk et al., 2022). Interestingly, self-organization of confined two dimensional (2D) iPSC colonies was hardly affected by YAP1 knockout as shown for the organization of TAZ and OCT4 within micro-contact printed iPSC colonies on day 6 (Figure 2E). Radial profile analysis (Figure 2F) and fluorescence microscopy (Figure 2G) showed an increased upregulation of NODAL at the border of *YAP*^*-/-*^ iPSC colonies compared to wildtype 2D colonies, which is in line with gene expression profiles from pluripotent wildtype and YAP1 deficient iPSCs. Overall, YAP1 does not seem to be a key regulator for self-organization of 2D pluripotent stem cell colonies.

### YAP1 deficiency enhances directed endodermal differentiation

Subsequently, we analyzed how YAP1 deficiency impacts directed differentiation toward endoderm, mesoderm, and ectoderm. Immunocytochemical analysis demonstrated a similar expression of the germ layer markers GATA6 (endoderm), brachyury (mesoderm), and PAX6 (ectoderm) in wildtype and *YAP*^*-/-*^ iPSCs (Suppl. Figure 2A). When we analyzed gene expression profiles of the differentiated cells (wildtype: n = 3, *YAP*^*-/-*^: n = 2), they clearly clustered according to the differentiation regimen in principal component analysis (PCA) (Figure 3A). Furthermore, endodermal differentiated YAP1 knockout samples clustered away from the respective wildtype cells. When we analyzed differential gene expression of *YAP*^*-/-*^ *versus* wildtype lines for each germ layer, the highest number of differential gene expression was observed for endoderm: 625 genes were downregulated and 420 genes were upregulated in YAP1 deficient iPSCs upon endodermal differentiation (log_2_ fold change > 1; adj. p-value < 0.05; Suppl. Figure 2B). Analysis of 10 canonical marker genes for each germ layer demonstrated an increased expression of endoderm related genes, whereas ectodermal marker genes were less expressed in *YAP*^*-/-*^ lines (Figure 3B). Even though changes in gene expression were not significant, tendencies are in line with previous reports from hESCs with YAP1 knockout (Hsu et al., 2018; Stronati et al., 2022).

**Figure 3:**
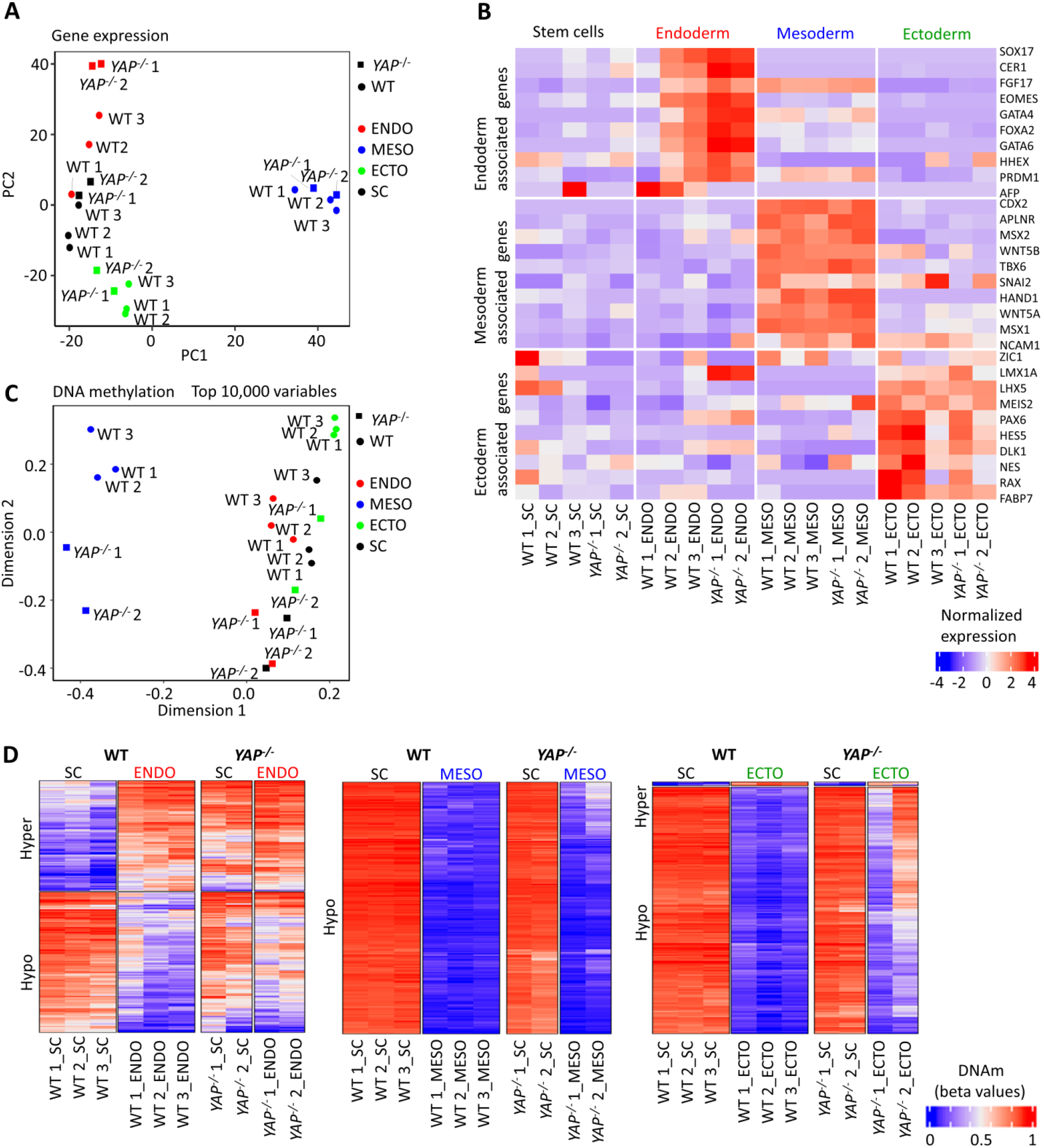
YAP1 deficiency enhances directed endodermal and decreases ectodermal differentiation. **(A)** Principal component analysis (PCA) based on gene expression profiles of WT (n = 3) and *YAP*^*-/-*^ iPSCs (n = 2) differentiated towards mesoderm, endoderm, and ectoderm by use of specific differentiation media, and undifferentiated control cells. **(B)** Normalized expression of ten canonical marker genes for each of the three germ layers in undifferentiated and differentiated WT and *YAP*^*-/-*^ iPSCs. **(C)** Multidimensional scaling (MDS) analysis of top 10,000 variable CpGs based on Illumina Beadchip Epic Array of undifferentiated and differentiated WT (n = 3) and *YAP*^*-/-*^ (n = 2) iPSCs. **(D)** DNAm levels (beta values) of top 200 significant CpGs with highest absolute difference in mean DNAm during differentiation of WT iPSCs toward endoderm, mesoderm, and ectoderm. DNAm levels are shown for undifferentiated and differentiated WT and *YAP*^*-/-*^ iPSCs

Next, we investigated if enhanced endodermal and reduced ectodermal differentiation of *YAP*^*-/-*^ lines were also reflected by epigenetic changes during directed differentiation. Overall, the DNAm profiles (wildtype: n = 3, *YAP*^*-/-*^: n = 2) clustered according to germ layer differentiation in multidimensional scaling (MDS). Particularly mesoderm was separated by the first dimension, whereas wildtype and *YAP*^*-/-*^ iPSCs were separated in the second dimension (Figure 3C). When we analyzed the difference in mean in DNAm between *YAP*^*-/-*^ and wildtype for each germ layer differentiation (adj. p-value ≤ 0.05; difference in mean ≥ 0.1), we found the strongest differences in hyper- and hypomethylated CpGs after mesodermal differentiation: 32,051 CpG sites were hypomethylated and 3,340 CpGs were hypermethylated in *YAP*^*-/-*^ lines (Suppl. Figure 2C). To analyze whether we find changes in DNAm at germ layer specific sites, we selected the top 200 significant CpGs with highest absolute difference in mean DNAm during differentiation of wildtype iPSCs toward endoderm, mesoderm, and ectoderm (Figure 3D). The DNAm changes of mesodermal differentiation were very similar in wildtype and *YAP*^*-/-*^ cells. Notably, for endodermal associated CpGs, we observed that the *YAP*^*-/-*^ iPSCs revealed such germ layer associated DNAm patterns already before differentiation. In contrast, the ectodermal associated DNAm changes of wildtype were less pronounced upon directed differentiation of *YAP*^*-/-*^ cells (Figure 3D). Differences in DNAm were more pronounced in *YAP*^*-/-*^ clone 2. DNAm as well as gene expression reflect changes in endoderm and ectoderm upon directed differentiation of YAP1 deficient iPSCs.

### YAP1 is required for early differentiation toward germ layers in embryoid bodies

While YAP1 deficiency was not responsible for 2D self-organization but clearly affected response of iPSCs to directed differentiation regimen, it was unclear if such lineage bias would also be observed in self-organized differentiation of embryoid bodies (EBs). We therefore generated 3D iPSC aggregates that were allowed to differentiate for five days in suspension culture. Viability of EBs was not reduced due to YAP1 knockout (Suppl. Figure 3A). Immunocytochemical analysis demonstrated that EBs generated from wildtype iPSCs express YAP1, downregulate the pluripotency marker OCT4, and upregulate compartmented expression of the germ layer markers GATA6, brachyury, and PAX6 (Figure 4A). In contrast, EBs from YAP1 deficient iPSCs maintained high expression of OCT4 and did not upregulate the representative germ layer markers, indicating that YAP1 is essential to trigger the self-organized multilineage differentiation in early iPSC aggregates (Figure 4B). To further confirm this finding, we artificially overexpressed *YAP1* with lentiviral transfection in the *YAP*^*-/-*^ lines (*YAP* OE). The restored YAP1 expression was demonstrated with Western Blot analysis (Suppl. Figure 3B) and immunofluorescence staining (Suppl. Figure 3C). Notably, artificial overexpression resulted in a predominant cytoplasmic localization, whereas in wildtype iPSCs YAP1 was rather located in the nucleus. Nevertheless, *YAP*^*-/-*^ EBs with artificial YAP1 overexpression restored even distribution of YAP1 expression and clear exit from pluripotent stage, as observed by downregulation of OCT4 expression. Furthermore, GATA6 and PAX6 were upregulated again with YAP1 overexpression, whereas the decrease of brachyury was not rescued (Figure 4C; Suppl. Figure 3C). These results were further substantiated on gene expression level with semi-quantitative RT-PCR (Figure 4D). Thus, YAP1 expression is essential for the self-organized 3D differentiation in early iPSC aggregates.

**Figure 4:**
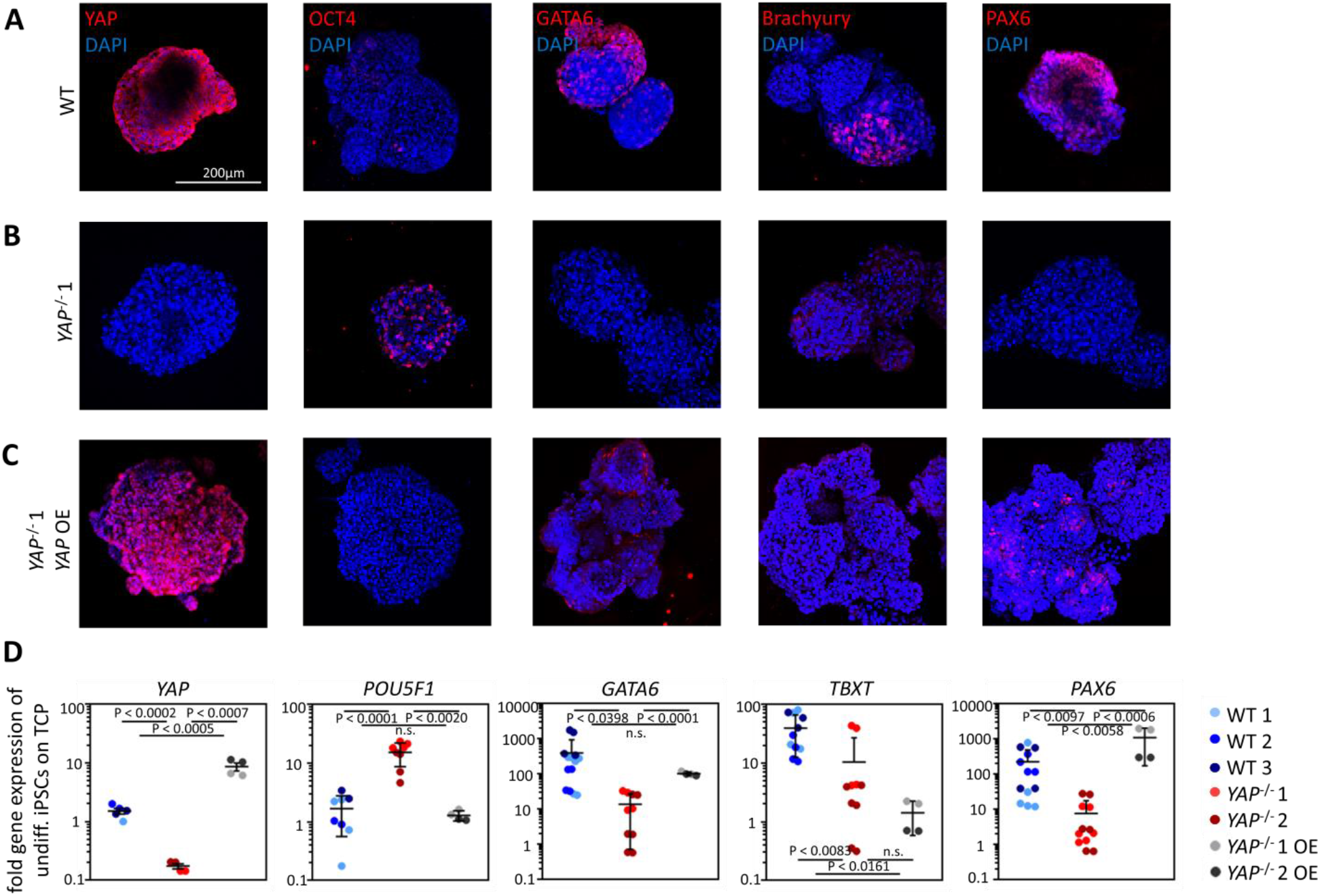
YAP1 is required for early germ layer differentiation in embryoid bodies. **(A)** Wildtype (WT) suspension embryoid bodies (EBs) were differentiated for five days in serum-containing medium. EBs were stained for either YAP1, OCT4, GATA6, brachyury, or PAX6 (nuclei counterstained with DAPI). In analogy, **(B)** *YAP*^*-/-*^ 1 EBs, **(C)** and *YAP1*^*-/-*^ EBs with constitutive overexpressing of YAP1 (*YAP*^*-/-*^ OE) were stained for the same markers. Imaging was performed via two-photon microscopy and representative images are depicted. **(D)** RT-qPCR analysis of *YAP1* expression, pluripotency (*POU5F1*) and germ layer-specific (*GATA6, TBXT*, or *PAX6*) markers for WT EBs, *YAP*^*-/-*^ EBs, and *YAP*^*-/-*^ *OE* EBs (WT and *YAP*^*-/-*^: n = 3 different experiments, *YAP* OE: n = 2 different experiments). Fold gene expression of undifferentiated iPSCs on tissue culture plastic (TCP)±SD; unpaired t-test. P-values > 0.05 are considered as not significant (n.s.)

### YAP1 modulates TGF-β signaling in non-directed differentiation

To further explore why YAP1 knockout lines were not capable for non-directed multilineage differentiation, we compared the transcriptome of *YAP*^*-/-*^ (n = 2) and wildtype (n = 3) EBs. We ended up with 176 genes that were significantly downregulated and 230 genes that were significantly upregulated in *YAP*^*-/*-^ EBs (log_2_ fold change > 1; adj. p-value < 0.05; Figure 5A). Again, *NODAL* (adj. p-value < 0.04, log_2_ fold change = 2.62) as well as its co-receptor teratocarcinoma-derived growth factor 1 (*TDGF1*; adj. p-value < 9.45*10^−9^, log_2_ fold change = 5.45), and its antagonist *LEFTY1 (*adj. p-value < 0.0006, log_2_ fold change = 6.93) were significantly upregulated in *YAP*^*-/-*^ EBs (Figure 5A). The downregulated genes were significantly enriched in functional categories for germ layer formation in gene set enrichment analysis (Figure 5B). Furthermore, expression of canonical marker genes for pluripotency remained higher expressed in YAP1 knockout EBs, whereas markers for the three germ layers where hardly upregulated (Figure 5C).

**Figure 5:**
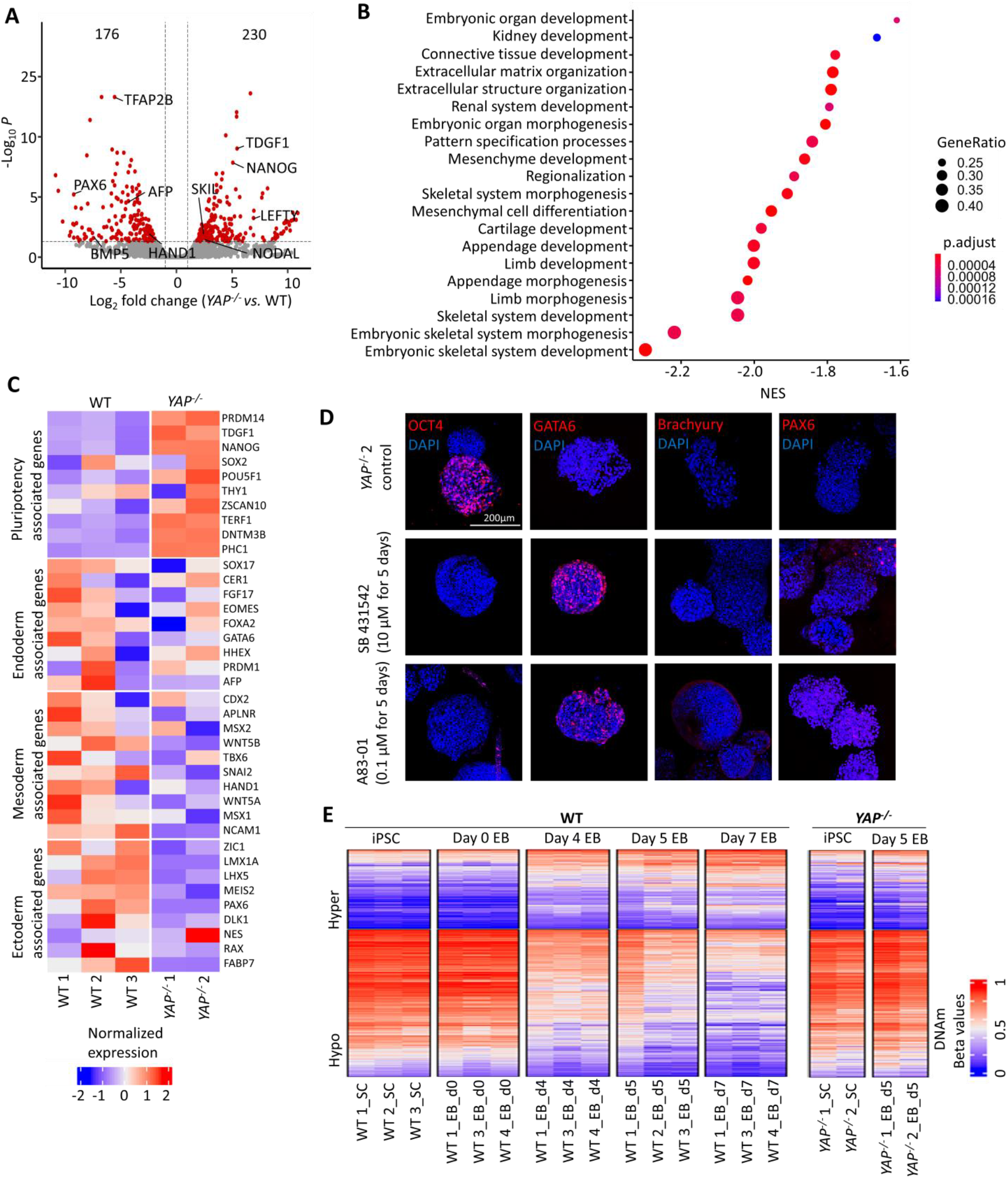
YAP1 modulates TGF-β signaling in non-directed differentiation. **(A)** RNA sequencing analysis of EBs from *YAP*^*-/-*^ (n = 2) and WT (n = 3) iPSCs. 176 genes are significantly downregulated and 230 genes are significantly upregulated in *YAP*^*-/-*^ (log_2_ fold change > 1; adj. p-value < 0.05). Specific genes related to germ layer formation or TGF-β and NODAL signaling are marked. **(B)** Gene set enrichment analysis of RNA sequencing data of EBs from *YAP*^*-/-*^ (n = 2) and wildtype (n = 3) iPSCs demonstrates significant differences in gene expression of early embryo development related categories. **(C)** Normalized expression of ten canonical marker genes for each of the three germ layers and pluripotent cells in wildtype and *YAP*^*-/-*^ EBs at day 5 of differentiation. **(D)** *YAP*^*-/-*^ EBs were generated for five days in medium without TGF-β inhibitors, or supplemented with 10 μM SB 431542 or 0.1 μM A83-01. Staining for OCT4, GATA6, brachyury, and PAX6 demonstrated that GATA6 and PAX6 expression could be restored by TGF-β inhibition. **(E)** Heatmap of top 1,000 significant CpGs (hypermethylated or hypomethylated) during wildtype EB differentiation from day 0 to day 4, and day 7. In these CpGs the EBs of *YAP*^*-/-*^ iPSCs reveal a similar pattern as non-differentiated iPSCs.

The functional relevance of NODAL signaling for germ layer formation in EBs was then further addressed with TGF-β type I-receptor inhibitors SB 431542 (10 μM) and A83 01 (0.1 μM) to our *YAP*^*-/-*^ EBs, respectively (Figure 5D). After five days of incubation with these inhibitors, the EBs were analyzed with immunofluorescence. OCT4 expression, which remained highly expressed in the untreated YAP1 knockout EBs, was downregulated upon TGF-β inhibitor treatment. However, this effect was more pronounced in the *YAP*^*-/-*^ 2 cell line as compared to the *YAP*^*-/-*^ 1 cell line. Furthermore, GATA6 upregulation was rescued by TGF-β inhibitor treatment in the *YAP*^*-/-*^ 2 cell line and both knockout lines revealed moderate upregulation of brachyury and PAX6 upon TGF-β inhibition.

Further evidence for an impaired differentiation was also observed on DNAm level. When we focused on the top 1,000 significant CpGs (either hypermethylated or hypomethylated) during seven days of wildtype embryoid body differentiation, we observed overall continuous DNAm changes. In contrast, *YAP*^*-/-*^ EBs at day 5, revealed very similar DNAm patterns as undifferentiated iPSCs (Figure 5E). No epigenetic changes could be observed for germ layer specific CpGs. Here, the top 200 CpGs with highest absolute difference in mean in DNAm during differentiation of wildtype iPSCs toward endoderm, mesoderm, or ectoderm (in analogy to Figure 3D) were analyzed in wildtype and *YAP*^*-/-*^ day 5 EBs. Similar to the 2D differentiation, endoderm related CpGs were hypermethylated in *YAP*^*-/-*^ EBs (Suppl. Figure 4A). Mesoderm and ectoderm related CpGs were hardly affected by EB formation of *YAP*^*-/-*^ cells, whereas some hypomethylation was observed in wildtype EBs (Suppl. Figure 4B,C). Consequently, impaired germ layer formation in self-organized *YAP*^*-/-*^ EBs is also clearly affected on DNAm level. Our results indicate that YAP1 is crucial to modulate the TGF-β pathway and to thereby orchestrate non-directed differentiation towards the germ layers.

## Discussion

The mechanotransducer YAP1 affects growth and differentiation of pluripotent cells (Dupont et al., 2011; Halder et al., 2012), but its specific functions have been controversially discussed. In this study, we generated human YAP1 knockout iPSC lines (*YAP*^*-/-*^) that recapitulate some features of previously described human YAP1 knockout ESC lines (Qin et al., 2016). *YAP*^*-/-*^ iPSCs showed colony formation and upregulation of the pluripotency marker OCT4. Furthermore, the Epi-Pluri-Score remained positive in *YAP*^*-/-*^ lines. In contrast, other authors suggested that YAP1 maintains iPSCs in naïve state and inhibits differentiation towards mesoderm lineages (Estarás et al., 2017; Lian et al., 2010; Qin et al., 2016). It has also been suggested that YAP1 rather stimulates differentiation towards ectoderm (Heng et al., 2020; Stronati et al., 2022). In fact, we observed significant differences on gene expression level between undifferentiated wildtype and *YAP*^*-/-*^ iPSCs and mesoderm associated genes were upregulated in *YAP*^*-/-*^ iPSCs, which might underline the before mentioned inhibition towards mesoderm lineages via YAP1 (Estarás et al., 2017). We demonstrated that *NODAL* and the related TGF-β and WNT pathways were differentially activated, which is in line with previous reports showing that YAP1 strongly regulates the activity of WNT and ACTIVIN signaling in pluripotent stem cells (Estarás et al., 2017; Hsu et al., 2018; Qin et al., 2016). Overall, our findings support the notion that YAP1, which is predominantly localized in the nucleus of iPSC colonies (Abagnale et al., 2017), acts as a co-transcription factor that modulates important signaling pathways, including NODAL, already at pluripotent state.

Relatively little is known on how YAP1 is involved in modulation of epigenetic changes, which ultimately determine cell fate decisions. We have directly compared DNAm levels of wildtype and *YAP*^*-/-*^ clones and came up with a number of differentially methylated CpGs in YAP1 deficient iPSCs – which might be unexpected given the phenotypic similarities. Yet, the differential DNAm changes were not directly related to genes with YAP1 binding sites, indicating that these specific epigenetic changes are not directly evoked by YAP1 transcriptional activity in human iPSCs. A recent study indicated that transient suppression of YAP1 in mouse ESCs might affect DNAm during four days of EB formation, possibly due to defective *de novo* methylation by down-regulation of Dnmt3l (Passaro et al., 2021). While we did not observe significant differences in gene expression of DNMT3L or other methyltransferases, we have also observed a higher number of CpGs with significant hypo-than hypermethylation in *YAP*^*-/-*^ clones. Overall, the distinct and reproducible epigenetic modifications seem to be governed by a mechanism that remains to be elucidated.

Colonies of iPSCs are heterogeneous – even under pluripotent culture conditions (Warmflash et al., 2014). We have recently demonstrated that particularly *NODAL* and its inhibitor *LEFTY* were amongst the most highly upregulated genes at the rim of spatially confined colonies (Elsafi Mabrouk et al., 2022). The interplay of NODAL and LEFTY was suggested to play a crucial role for the pattern development in reaction-diffusion models (Etoc et al., 2016; Müller et al., 2012; Tewary et al., 2017). Here, we demonstrate that despite pronounced upregulation of NODAL the self-organization of 2D YAP1 knockout colonies was not affected. Furthermore, it has been suggested that YAP1 is directly involved in the recognition of topographic stimuli which induce proliferation and differentiation of iPSCs (Raghunathan et al., 2014; Reimer et al., 2016). Our previous work additionally pointed to a potential relevance of YAP1 for the elongation of individual iPSCs and iPSC colonies along sub-micrometer structures (Abagnale et al., 2017). Such structures guide the organization of iPSC colonies by directing apical actin fibers and cell division planes and there was evidence that YAP1 might be involved in this process (Abagnale et al., 2017). However, our results demonstrate that YAP1 is not essential to mediate this cellular response to sub-micrometer surface topography. It is conceivable that the paralog of YAP1 – TAZ (WWTR1) – counteracts some of the effects of YAP1 deficiency (Plouffe et al., 2018).

While YAP1 is dispensable for maintenance of pluripotency and self-organization of pluripotent colonies, it is clearly involved in the regulation of differentiation. Directed differentiation towards endoderm, mesoderm, and ectoderm could still be achieved with *YAP*^*-/-*^clones, while there was evidence for modification of endoderm and ectoderm differentiation in YAP1 knockout. These findings are also in line with a recent study on hESCs demonstrating that the self-organized differentiation of *YAP*^*-/-*^ hESCs in colonies display reduced ectodermal and enlarged mesodermal and endodermal areas (Stronati et al., 2022). Controversially to previous publications (Estarás et al., 2017; Stronati et al., 2022), YAP1 deficiency did not modulate mesodermal differentiation on gene expression level. Interestingly, our non-directed 3D multi-lineage differentiation in embryoid bodies clearly demonstrated that cell-fate decisions toward endoderm, mesoderm, and ectoderm are severely impaired in YAP1 knockout iPSCs. The significant upregulation of NODAL in the *YAP*^*-/-*^ EBs was apparently not sufficient to drive endodermal and mesodermal differentiation, as previously observed in a 2D “gastruloid model” (Stronati et al., 2022). On the other hand, the phenotype of differentiated *YAP*^*-/-*^ EBs could at least partially be rescued by constitutive re-expression of *YAP1* as well as by inhibition of the TGF-β type I-receptor. Taken together, in early iPSC aggregates YAP1 seems to control the TGF-β pathway and thereby self-organized germ layer specification.

## Conclusion

While cultured under pluripotent culture conditions YAP1 knockout iPSCs reveal a relatively normal phenotype in growth and self-organization of 2D colonies, albeit gene expression changes – particularly up-regulation of *NODAL* – and epigenetic modifications already point to modification of germ layer differentiation. In line with other studies, YAP1 deficiency leads to changes in endodermal and ectodermal differentiation potential (Heng et al., 2020; Stronati et al., 2022). Interestingly, while directional differentiation was still possible in *YAP1*^*-/-*^ cells, germ layer development was almost completely impaired in non-directional differentiation in embryoid bodies. The essential role for the self-organization process mediated by YAP1 seems to be governed by modulation of TGF-β signaling. The different mechanobiological context of cellular differentiation may contribute to the multilayered functions of YAP1.

## Author Contributions

K.Z. and R.G. generated *YAP*^*-/-*^ iPSC lines. K.Z. performed experiments, analyses of cell-based experiments, and generated samples for RNA sequencing and DNA methylation analyses. M.E.M. performed gene expression analyses and confocal microscopy. M.S. performed analyses of DNAm profiles. C.M. generated *YAP1* overexpressing cell lines. K.Z. and A-C.H. performed two-photon microscopy. C.H. and A.G. generated structured polyimide foils. K.Z., R.G., M.Z., and W.W. designed and supervised the study. K.Z. and W.W. wrote the manuscript and all authors approved the final version.

## Acknowledgements

This research project was supported by the Core Facilities “Two-Photon Imaging” and “Confocal Microscopy”, Core Facilities of the Interdisciplinary Center for Clinical Research (IZKF) Aachen within the Faculty of Medicine, RWTH Aachen University; by the Federal Ministry of Education and Research (GO-Bio: Pluri-Screen, 16LW0017); the Deutsche Forschungsgemeinschaft (DFG, German Research Foundation - 363055819/GRK2415; WA1706/11-1; WA 1706/12-1 within CRU344; WA1706/14-1); by the START-Program of the Faculty of Medicine, RWTH Aachen (01/20); and the ForTra gGmbH für Forschungstransfer der Else Kröner-Fresenius-Stiftung.

