## Supplemental Figures and Tables for "YAP1 is essential for self-organized differentiation of pluripotent stem cells"

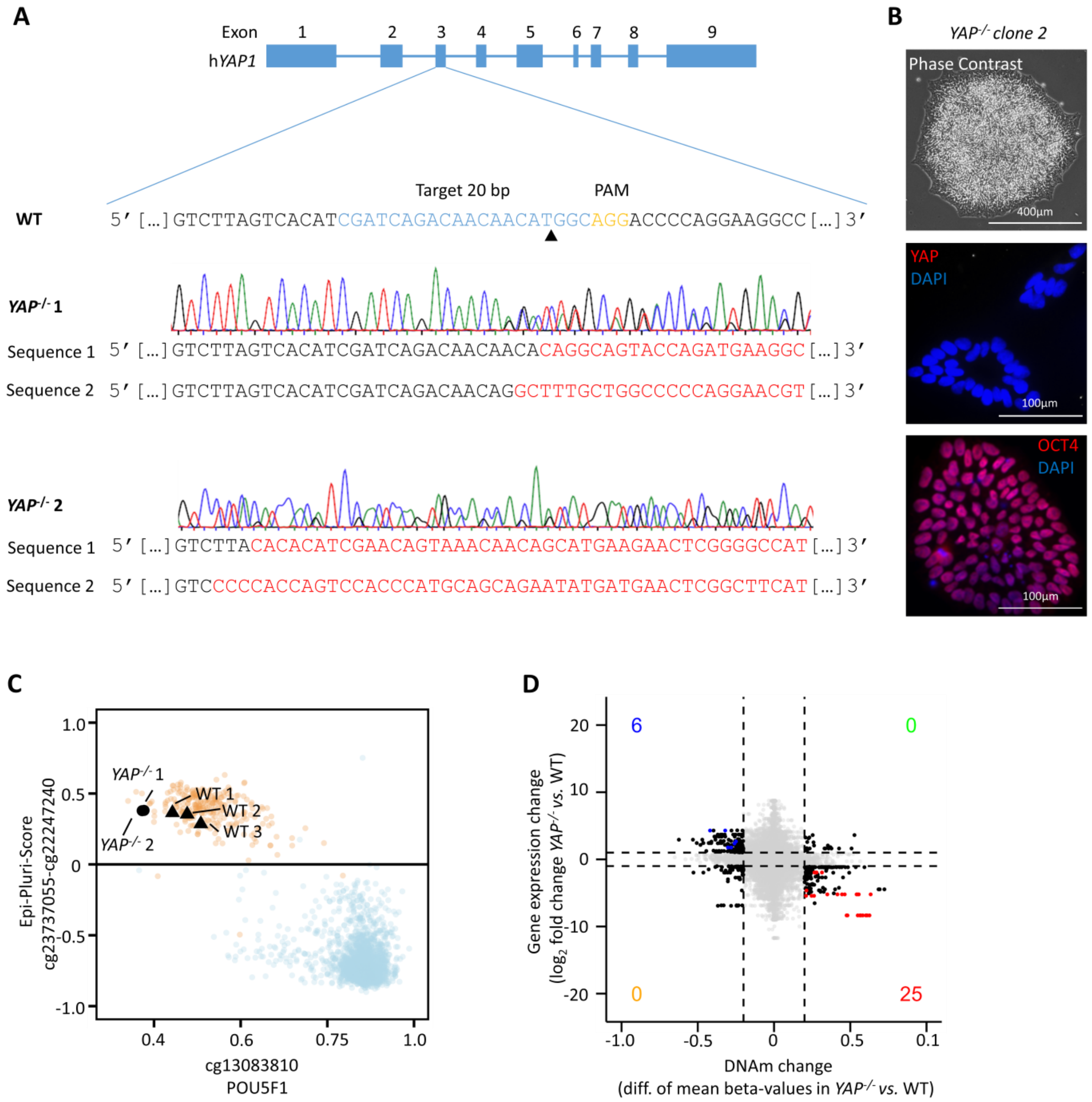


**Supplemental Figure 1: Generation of *YAP*$^{-/-}$ iPSCs with CRISPR/Cas9**
**(A)** Two clonal human YAP1 knockout iPSCs lines (*YAP*$^{-/-}$ 1 and *YAP*$^{-/-}$ 2) were generated with homozygous frame shift mutations in exon three by CRISPR/Cas9 technology. Gene sequences were analyzed with Sanger-sequencing. **(B)** Exemplary depiction of a phase contrast image and immunofluorescence analysis of YAP1, OCT4, and DAPI for the *YAP*$^{-/-}$ 2 iPSC line (in analogy to Figure 1B). **(C)** Epi-Pluri-Score analysis of wildtype (WT; n = 3) and *YAP*$^{-/-}$ (n = 2) iPSC lines validates pluripotent state. The assay is based on DNAm at three specific CpGs: a CpG in *POU5F1* (cg13083810), and two CpGs in *ANKRD46* and *C14orf115* (cg23737055 and cg 22247240), which are combined as Epi-Pluri-Score (Lenz et al., 2015). A positive Epi-Pluri-Score is indicative for pluripotency. The reference clouds refer to DNAm profiles (all Illumina HumanMethylation27 BeadChip platform) of 264 pluripotent (orange) and 1,951 non-pluripotent cell preparations (blue). **(D)** Association of DNA methylation changes and corresponding gene expression changes in WT *versus YAP*$^{-/-}$ iPSCs. Only CpGs in promoter regions (TSS1500, TSS200) are considered. Each dot represents a gene-CpG-pair (genes as well as CpGs might be duplicated). Colored dots depict pairs with a significant difference in DNAm (adj. p-value ≤ 0.05; difference in mean ≥ 0.1) and significant gene expression changes ($log_2$ fold change > 1; adj. p-value < 0.05). In tendency, genes with hypomethylation in promoter regions are upregulated on gene expression level, and *vice versa*.

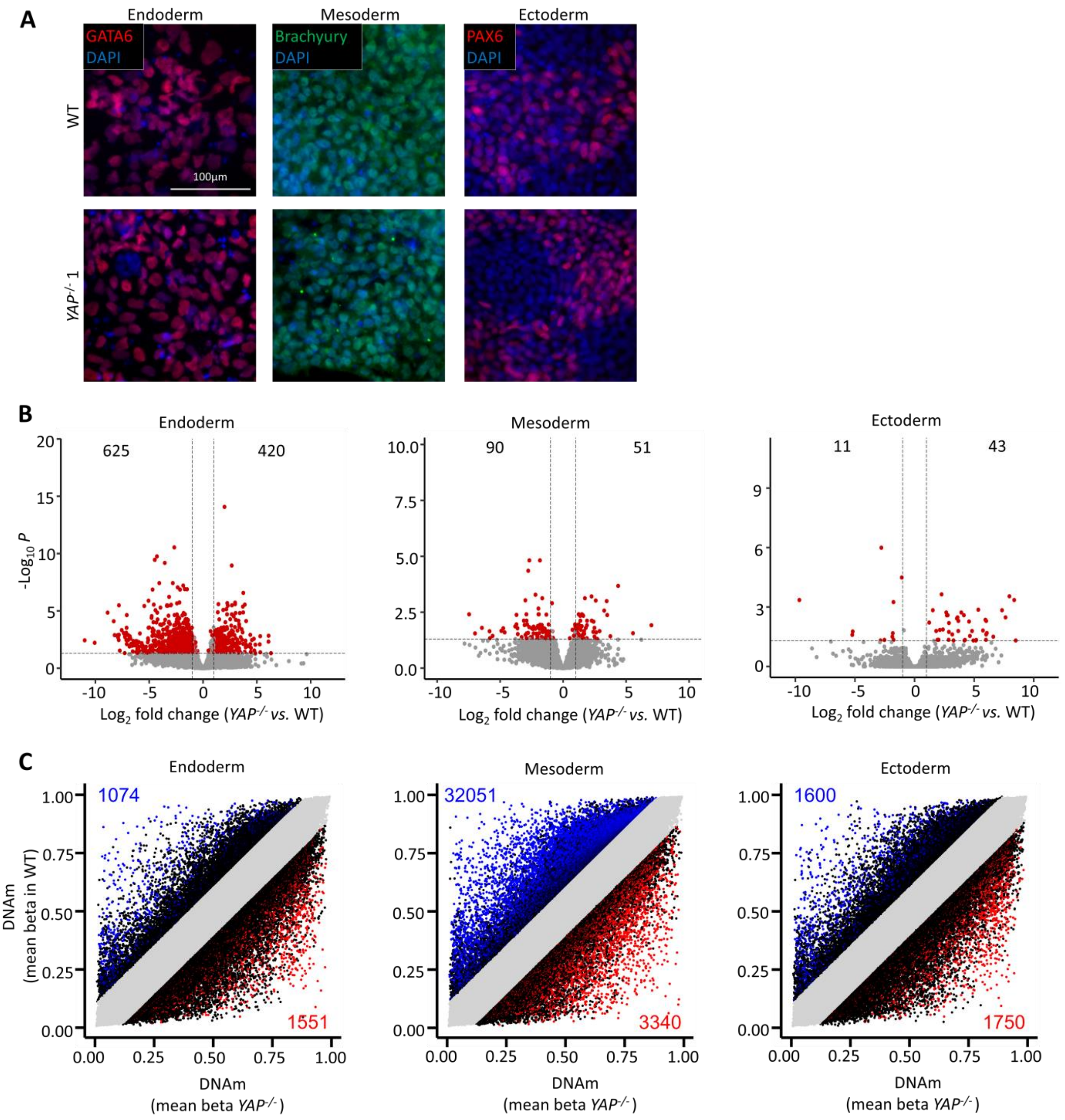


**Supplemental Figure 2: *YAP1$^{-/-}$* iPSCs are capable of directed trilineage differentiation**
**(A)** Exemplary immunofluorescence images of WT and *YAP$^{-/-}$* iPSCs upon directed differentiation toward endoderm, mesoderm, and ectoderm. The corresponding marker genes were also upregulated in *YAP$^{-/-}$* cells (GATA6, brachyury, and PAX6, respectively, nuclei counterstained with DAPI). **(B)** Gene expression changes between *YAP$^{-/-}$* and WT iPSCs (RNA sequencing analysis; n = 2 and n = 3, respectively) after directed endodermal, mesodermal, and ectodermal differentiation, respectively. Upon endoderm differentiation the differences were most pronounced with upregulation of 420 and downregulation of 625 genes in *YAP$^{-/-}$* iPSCs ($\log_2$ fold change > 1; adj. p-value < 0.05). **(C)** Scatter plot of global DNA methylation changes between *YAP$^{-/-}$* (n = 2) and WT (n = 3) iPSCs after endodermal, mesodermal, and ectodermal differentiation. DNAm was analyzed with Illumina EPIC BeadChips (mean beta-values are depicted). In mesodermal differentiated cells, most CpGs reveal significant differences between *YAP$^{-/-}$* and WT (hypomethylation of 32,051, hypermethylation of 3,340 CpGs in *YAP$^{-/-}$* iPSCs; adj. p-value ≤ 0.05; difference in mean beta value ≥ 0.1).

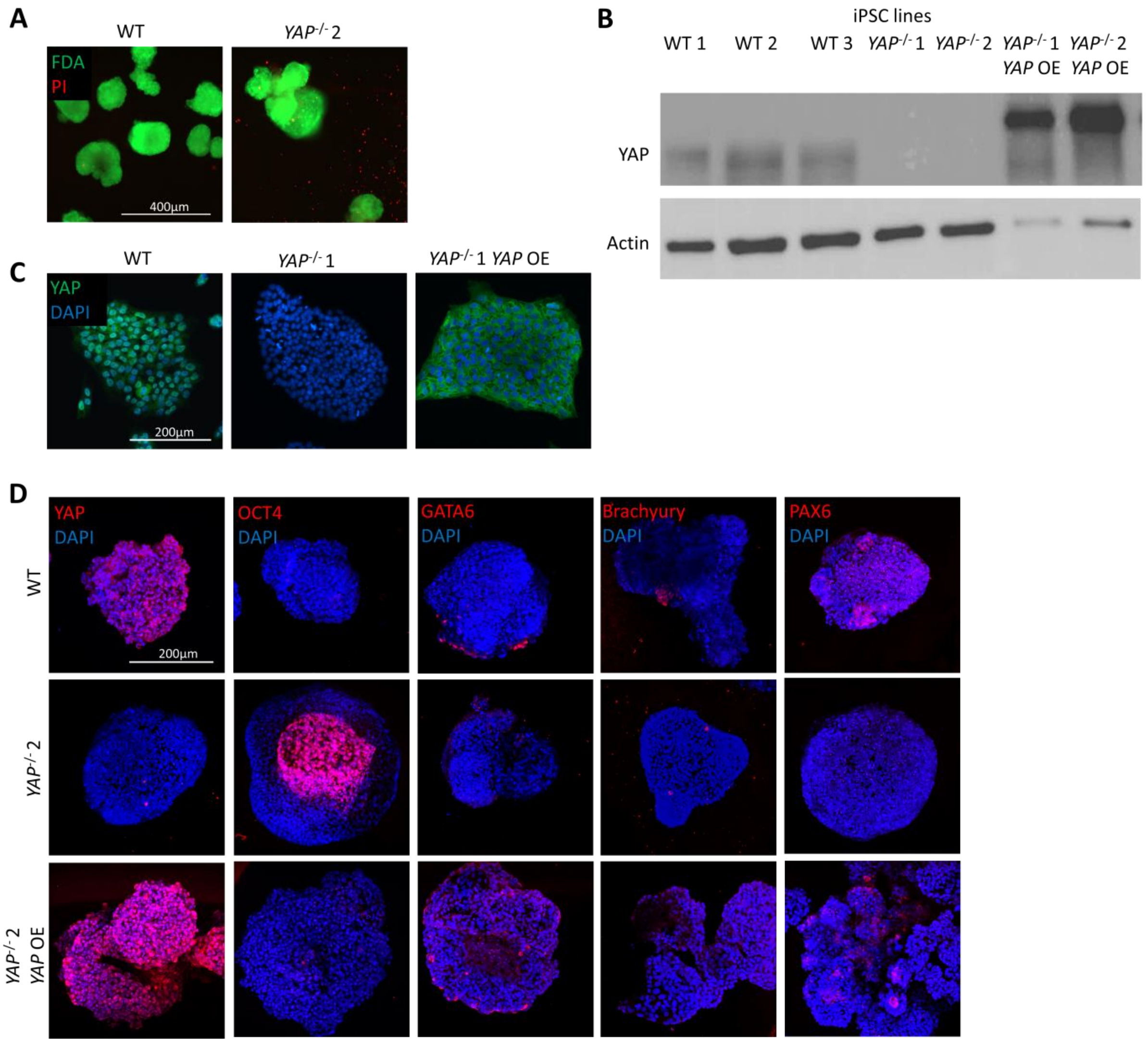


**Supplemental Figure 3: YAP1 overexpression can partially rescue *YAP*$^{-/-}$ phenotype**

**(A)** Live/dead staining of embryoid bodies from WT and *YAP*$^{-/-}$ iPSCs at day 6 of differentiation in serum-containing medium. Viable cells are stained with fluorescein diacetate (FDA; green), dead cells are stained with propidium iodide (PI; red). **(B)** Western Blot analysis of three WT, two *YAP*$^{-/-}$, and two *YAP*$^{-/-}$ iPSC lines with constitutive YAP1 overexpression (YAP OE) for antibodies against YAP1 (65 kD) and the housekeeping protein actin (42 kD). For YAP OE samples only 10 µg proteins were used. **(C)** Immunofluorescence analysis of YAP1 expression in iPSC colonies of WT, *YAP*$^{-/-}$ 1, and *YAP*$^{-/-}$ *1 OE* clones (nuclei counterstained with DAPI). YAP1 overexpression is homogeneous in *YAP*$^{-/-}$ *1 OE* but also in the cytoplasm. **(D)** Embryoid bodies were generated from WT, *YAP*$^{-/-}$ 2, and *YAP*$^{-/-}$ 2 OE clones for five days in serum-containing medium. EBs were stained for either YAP1, OCT4, GATA6, brachyury, or PAX6 (nuclei counterstained with DAPI).

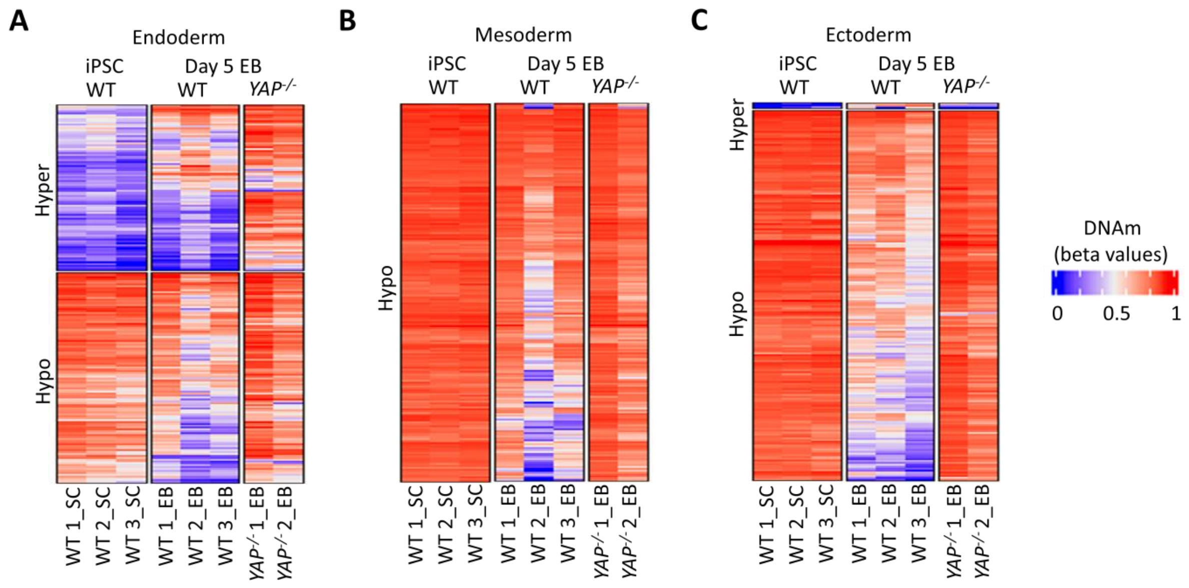


**Supplemental Figure 4: Germ layer associated DNAm changes in *YAP*$^{-/-}$ EBs**

DNA methylation levels (beta values) of top 200 significant CpGs with highest absolute difference in mean in DNAm during differentiation of WT iPSCs toward **(A)** endoderm, **(B)** mesoderm, and **(C)** ectoderm. All of these signatures show clear changes in the directed differentiation (as depicted in Figure 3D). Here, DNAm levels of the same CpGs are shown for undifferentiated WT iPSCs, EBs from WT iPSCs 5 days after differentiation, and *YAP*$^{-/-}$ EBs 5 days after differentiation.

### Supplemental tables

**Table S1: Antibody list**

| | NAME | CLONE | COMPANY | DILUTION |
|---|---|---|---|---|
| WB | YAP | EP1674Y | Abcam, Cambridge, UK | 1:5,000 |
| | Beta-Actin | Ac-74 | Sigma-Aldrich, St. Louis,USA | 1:10,000 |
| | Goat IgG anti-rabbit IgG-HRPO | polyclonal | Dianova, Hamburg, Germany | 1:10,000 |
| | Goat IgG anti-mouse IgG-HRPO | polyclonal | Dianova, Hamburg, Germany | 1.10,000 |
| Immunofluorescence | YAP | 63.7 | Santa Cruz, Dallas, USA | 1:50 |
| | TAZ | polyclonal | Abcam, Cambridge, UK | 1:100 |
| | OCT4 | polyclonal | Abcam, Cambridge, UK | 1:500 |
| | GATA6 | D61E4 | Cell Signaling, Danvers, USA | 1:1,600 |
| | Brachyury | polyclonal | R&D Systems, Minneapolis, USA | 1:20 |
| | PAX6 | AD2.35 | Santa Cruz, Dallas, USA | 1:200 |
| | NODAL | 5C3 | Sigma-Aldrich, St. Louis, USA | 1:100 |
| | Donkey anti-goat (Alexa Fluor 488) | | Invitrogen, Waltham, USA | 1:200 |
| | Donkey anti-goat (Alexa Fluor 594) | | Invitrogen, Waltham, USA | 1:200 |
| | Goat anti-rabbit (Alexa Fluor 488) | | Invitrogen, Waltham, USA | 1:200 |
| | Goat anti-rabbit (Alexa Fluor 594) | | Invitrogen, Waltham, USA | 1:200 |
| | Goat anti-mouse (Alexa Fluor 488) | | Invitrogen, Waltham, USA | 1:200 |
| | Goat anti-mouse (Alexa Fluor 594) | | Invitrogen, Waltham, USA | 1:200 |

Listed are primary and secondary antibodies that were used for Western blot (WB) and immunofluorescence analysis.

**Table S2: Primer list**

| | GENE | NAME | DNA SEQUENCE |
|---|---|---|---|
| RT-qPCR primer | *POU5F1* | POU5F1 For | GGGGGTTCTATTTGGGAAGGTA |
| | *POU5F1* | POU5F1 Rev | ACCCACTTCTGCAGCAAGGG |
| | *GATA6* | GATA6 For | CTCAGTTCCTACGCTTCGCAT |
| | *GATA6* | GATA6 Rev | GTCGAGGTCAGTGAACAGCA |
| | *TBXT* | Brachyury For | CAGTGGCAGTCTCAGGTTAAGAAGGA |
| | *TBXT* | Brachyury Rev | CGCTACTGCAGGTGTGAGCAA |
| | *PAX6* | PAX6 For | TCGAAGGGCCAAATGGAGAAGAGAAG |
| | *PAX6* | PAX6 Rev | GGTGGGTTGTGGAATTGGTTGGTAGA |
| | *GAPDH* | GAPDH For | GAAGGTGAAGGTCGGAGTC |
| | *GAPDH* | GAPDH Rev | GAAGATGGTGATGGGATTTC |

Listed are forward (For) and reverse (Rev) RT-qPCR primer.

**Table S3: TaqMan gene expression assays**

| GENE | NAME | COMPANY |
|---|---|---|
| *YAP1* | Hs00902712_g1 YAP1 | Applied Biosystems, Waltham, USA |
| *GAPDH* | Hs02758991_g1 GAPDH | Applied Biosystems, Waltham, USA |

Listed are TaqMan gene expression assays for RT-qPCR.
